# Translatome-based classification reveals a dual metabolic dependency of a new tumor subtype of pancreatic cancer

**DOI:** 10.1101/2020.12.23.424227

**Authors:** Sauyeun Shin, Remy Nicolle, Christine Jean, Remi Samain, Mira Ayadi, Jerome Raffenne, Alexia Brunel, Jacobo Solorzano, Cindy Neuzillet, Carine Joffre, Stephane Rocchi, Juan Iovanna, Nelson Dusetti, Ola Larsson, Stephane Pyronnet, Corinne Bousquet, Yvan Martineau

## Abstract

Molecular profiling of Pancreatic Ductal Adenocarcinoma (PDA), based on transcriptomic analyses, identifies two main prognostic subtypes (basal-like and classical), but does not allow personalized first-line treatment. To date, tumors have not been profiled based on protein synthesis rates, yet the step of mRNA translation is highly deregulated in both PDA cancer cells and their microenvironment. Using a collection of twenty-seven pancreatic Patient-Derived Xenografts (PDX), we performed genome-wide analysis of translated mRNA (translatome). Unsupervised bioinformatics analysis revealed a new tumor subtype harboring a low protein synthesis rate, but associated with a robust translation of mRNAs encoding effectors of the integrated stress response (ISR), including the transcription factor ATF4. Functional characterization of the “ISR-activated” human cancer cells revealed a high resistance to drugs, low autophagic capacities, and importantly, metabolic impairments in the serine synthesis and transsulfuration pathways. Overall, our study highlights the strength of translatomic profiling on PDA, which here revealed an unforeseen drug-resistant cancer cell phenotype, whose auxotrophy to both serine and cysteine may be amenable to targeted therapy.

## INTRODUCTION

Pancreatic ductal adenocarcinoma (PDA) is a lethal disease with a 5-year survival rate below 10%, and represents the 4^th^ cause of cancer-related death in industrialized countries (Rahib et al. 2020). This poor prognosis is mainly due to late diagnosis (often at metastatic stages) and the lack of efficient first-line chemotherapies. Indeed, Gemcitabine/Abraxane and FOLFIRINOX, show low response rates (23 and 31%, respectively) in PDA patients (Narayanan and Weekes 2015).

“Omic” studies led to important characterization of PDA biology (Jones et al. 2008) as well as patient stratification (Moffitt et al. 2015). For example, exome sequencing revealed in a small group of patients (11%), mutations of DNA repair genes, which explain increased patient response to platinum-based chemotherapies (Waddell et al. 2015). Nonetheless, all the transcriptomic classifications of PDA tumors which have been reported today, agree on the existence of two main subtypes (basal-like and classical), which are prognostic but not predictive of sensitivity to a specific chemotherapy.

PDA tumor cells are characterized by a high plasticity, enabling their adaptation to the harsh PDA microenvironment (poor in oxygen and nutrients). Although stromal cells support survival of cancer cells, the latter have also developed high intrinsic abilities to survive stress conditions. Before any transcriptional response is elicited, cells first adapt by shutting down the protein synthesis, one of the most energy-consuming processes, and then by engaging a coordinated stress response (Pakos-Zebrucka et al. 2016). Protein synthesis was previously reported to be deregulated in PDA, supporting tumor growth and survival (Chio et al. 2016; Martineau et al. 2013). Accordingly, activation of signaling pathways, or altered expression of translation regulators, was reported in PDA to impact on mRNA translation and to favor adaptation to environmental changes (Morran et al. 2014; Goh et al. 2019; Wang et al. 2019; Müller et al. 2019).

Importantly, protein expression level is not strictly correlated to the mRNA abundance (Schwanhäusser et al. 2011), but principally relies on the translation efficiency (TE) of the mRNA. One of the best characterized examples is the translational induction of the activating transcription factor ATF4 mRNA, following activation of the Integrated Stress Response (ISR), encompassing phosphorylation of eIF2α and global protein synthesis attenuation. Consequently, the amount of ATF4 cannot be correlated to the abundance of its transcript (Pakos-Zebrucka et al. 2016). ATF4 is a short-lived transcription factor, critical for survival and adaptation to numerous cellular stresses, such as oxidative stress, amino acid starvation, misfolding of Endoplasmic Reticulum (ER)-located proteins, and chemotherapy exposure (Pakos-Zebrucka et al. 2016). Therefore, previous transcriptome-based PDA classifications have omitted an important parameter, which is the cellular mRNA translation activity. Thus, to test whether, beyond transcription, mRNA translation could improve PDA patient stratification, we performed a genome-wide analysis of highly translated mRNAs (translatome) using a collection of 27 pancreatic patient-derived xenografts (PDXs). Herein, we identified a subgroup of tumors with low protein synthesis rate, while whilst associated with a high translation efficiency of a subset of mRNAs implicated in ISR. Using primary PDX-derived cell cultures and commonly-used PDA tumor cells, we show that features such as high resistance to apoptosis, low autophagic capacity, and dependency on exogenous amino-acids, characterize this “ISR-activated” phenotype.

## RESULTS

### Translatome-based PDA subtyping

In order to define whether examination of mRNA Translation Efficiency (TE) can allow identifying novel PDA tumor subtypes, we performed translatomics on a collection of twenty-seven patient-derived xenografts (PDX), previously used to classify PDA tumors through transcriptomics (Nicolle et al. 2017). PDXs are considered as the most reliable avatars of individual patient tumors, hence recapitulating the histology (including stroma), the genetic and molecular heterogeneities of the primary tumor (Knudsen et al. 2018). PDXs are also a hybrid model where human cancer cells proliferate and remodel the murine stroma (the initial human stroma is lost with the first passage in mouse). Using Simultaneous MApping for Pdx algorithm (SMAP) (Nicolle et al. 2017), human tumor RNAs can be separated computationally from mouse stroma RNAs, in order to analyze transcriptional and translational activities specifically in human cancer cells within the PDX. The SMAP algorithm requires sufficiently long RNA sequences to ensure accurate reads alignment to distinguish mouse and human RNAs. Thus, Riboseq approaches, generating short sequences of only 35 nucleotides (Ingolia 2016), were not adapted to achieve reliable alignments to the distinct genomes, contrary to the traditional polysome-profiling which can be coupled to standard RNAseq generating longer reads. Consequently, polysome-profiling adapted to small tissue samples (based on discontinuous sucrose gradient, Liang et al. 2018), was used to purify actively translated mRNAs from tumors (Fig. 1A, Supp. Fig. 1A). The ratio of polysome-associated mRNA to total transcript level of each mRNA was obtained using Anota2seq-based estimation (Oertlin et al. 2019). Anota2seq also integrates changes in total mRNA levels, which could also impact on the pool of polysome-associated mRNA, allowing the identification of mRNAs with bona fide changes in TE. TE of each transcript in each tumor were quantified, followed by independent component analysis (ICA, Fig. 1A). After suppression of experimental and analytical bias, this strategy allowed identification of 4 independent components (IC). We focused on IC3, which showed enrichment with an ER stress translational signature, especially for the TE of prototypical mRNA such as ATF4, JUN and ATF5 (Supp. Fig. 1B), while it anti-correlated with TEs of mRNAs encoding olfactory receptors (olfactory transduction KEGG pathway; NES: −2.52, P-value: 9.9e-03). This multi-gene family, comprising around 800 genes, is over-represented in the genome. This suggests a global attenuation of protein synthesis in IC3 (Fig. 1B). Scatter plots of ATF4 polysomal vs. total mRNA for each PDX tumor, show high association between ATF4 mRNA TE and the IC3 component (Fig. 1C, left panels). Conversely, IC3 is not associated with the TE of mRNA encoding OR8G1, a member of the olfactory receptor family and one of the most translationally repressed gene in IC3 (Fig. 1C, right panels).

**Figure 1:**
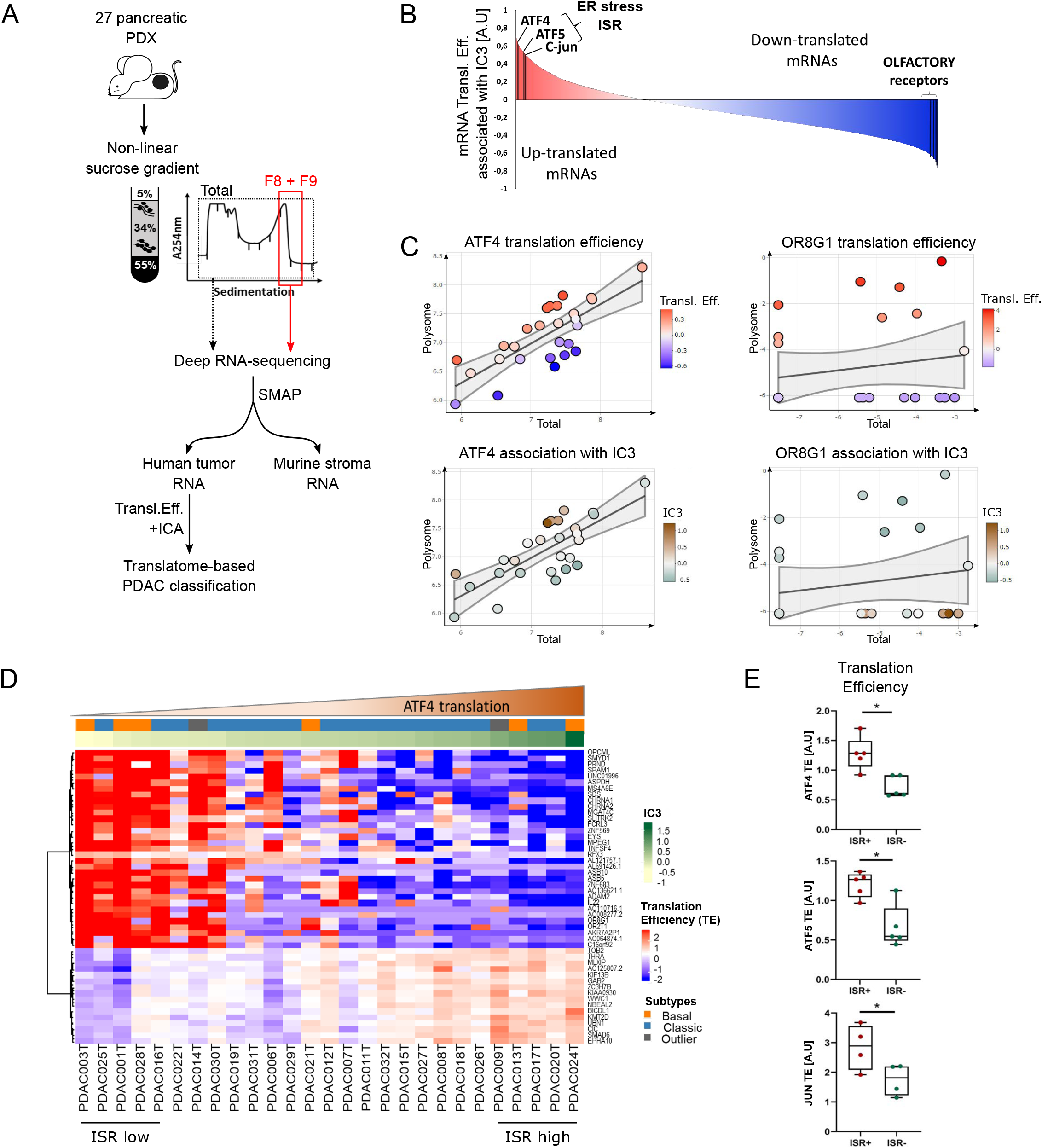
Translatomics-based PDA subtyping. **A)** Schematic of the translatome-based PDA classification procedure. PDX lysates were loaded on a non-linear sucrose gradient. Total and polysomal fraction (F8/F9) were purified and sequenced. SMAP algorithm was used to separate human and murine RNA sequences. The Anota-2seq package followed by Independent component analysis (ICA) allowed identification of components with a translational signature. **B)** Representation of mRNA Translation Efficiency (positive: red, negative: blue) associated with IC3. **C)** Scatter plot representing ATF4 (left) and OR8G1 (right) polysomal (*y axis*) and total mRNA (*x axis*) abundance in each PDX. Translation efficiency (TE) are shown in red/blue and association with IC3 in brown/green. **D)** Heat-map representing the top correlated gene translation efficiency (red, high TE/ blue, low TE) of PDX tumors associated with IC3 (green), and ATF4 expression (brown). Basal-like and classical classification of PDX are indicated in respectively orange and blue. **E)** The indicated mRNA TE have been plotted using the top 5 PDX correlated (ISR+) or not correlated (ISR-) with IC3. Data are presented as mean ± SD. P-values were calculated using unpaired t-test (*P< 0.05).

Translational regulation during Endoplasmic Reticulum (ER) stress is part of the ISR. ISR reduces cap-dependent translation initiation, mainly by impacting the ternary complex formation through the blockade of eIF2-GDP recycling. ISR is mainly mediated by phosphorylation of eIF2α by 4 different kinases, PERK (Protein kinase R-like endoplasmic reticulum kinase), GCN2 (General control nonderepressible 2), HRI (Heme-regulated inhibitor) or PKR (protein kinase R) upon ER stress, amino acid deprivation, heme deficiency or viral infection, respectively. This leads to a global reduction of protein synthesis, and also activates translation of specific mRNAs such as ATF4, ATF5 or c-JUN (Guan et al. 2017). The majority of IC3 top-correlated genes showed a low RNA TE, suggesting a global down-regulation of mRNA translation (Fig. 1D blue), contrary to a translational enhancement of a small subset of mRNAs (Fig. 1D red). Indeed, IC3 was associated with a decreased density of polysome mRNA counts (Supp. Fig. 1C). Importantly, IC3 did not correlate with the reported transcriptomic basal-like and classical pancreatic molecular subtypes (Fig. 1D orange, blue). The five tumors with the best (ISR-high), and the worst (ISR-low), association with IC3 were selected, and data corresponding to TE were plotted for ATF5, c-JUN and ATF4 mRNA (Fig 1E), which were further confirmed by RT-qPCR for ATF4 (Supp. Fig. 1D), highlighting the fact that the TE of mRNAs encoding these genes is significantly associated to the ISR-high tumors. Altogether, this study identifies a new tumor subset associated with the IC3, characterized by specific translational features linked to high ISR activation.

### Translation properties of ISR-activated cells

To characterize the IC3-associated PDA phenotype, two primary PDX-derived cancer cultures were isolated from one of each ISR-high (017T) and ISR-low (003T) tumors. Elevated levels of ATF4 expression and eIF2α phosphorylation were found in 017T cells, as compared to 003T cells, in accordance with the tumor subtype (Fig. 2A left, Supp. Fig. 2A left). A similar analysis, performed on commonly-used pancreatic cancer cell lines revealed that AsPC-1 cells show the highest expression of ATF4 in association with a higher proportion of phosphorylated eIF2α compared to the total eIF2α, as opposed to MiaPaca-2 or Panc-1 cells (Fig. 2A right, Supp. Fig. 2A right). Importantly, 017T and AsPC-1 cells had reduced amounts of total eIF2α, as compared to other cells, suggesting a higher propensity to ISR activation. Hereafter, AsPC-1 and 017T are referred to “ISR-activated” cells, whereas MiaPaca-2 and 003T cells to “translational reference” cells. In ISR-activated cells, ATF4 mRNA was not overexpressed, but showed an enhanced TE, as evidenced by an enrichment in heavy polysome fractions in linear sucrose gradient (Fig. 2B). Conversely, distribution of OR8G1 mRNA, a member of the over-abundant olfactory receptor family, showed enrichment in 003T heavy polysome fractions, whereas it was decreased in 017T (Supp. Fig. 2B), as observed in PDX samples (Fig. 1C). ISR-activated cells also showed reduced protein synthesis rate, as measured by [^35^S]-methionine incorporation and polysome abundance (Fig. 2C, D). To fully demonstrate the maintenance of ISR in 017T and AsPC-1 cells, ISRIB, an inhibitor of ISR, was used to counteract eIF2α phosphorylation (Zyryanova et al. 2018). As expected, ISRIB favored mRNA translation in ISR-activated cells whereas it had limited effect on MiaPaca-2 cells, as indicated by puromycin incorporation (Fig. 2E). In addition, in ISR-activated cells, ISRIB treatment reduced ATF4 abundance without impacting eIF2α phosphorylation (Fig. 2F). These data demonstrate that the ISR-activated phenotype uncovered in PDX (Fig. 1) is conserved from primary tumors to cell culture dishes, despite the unlimited supply of nutrients and oxygen in culture medium.

**Figure 2:**
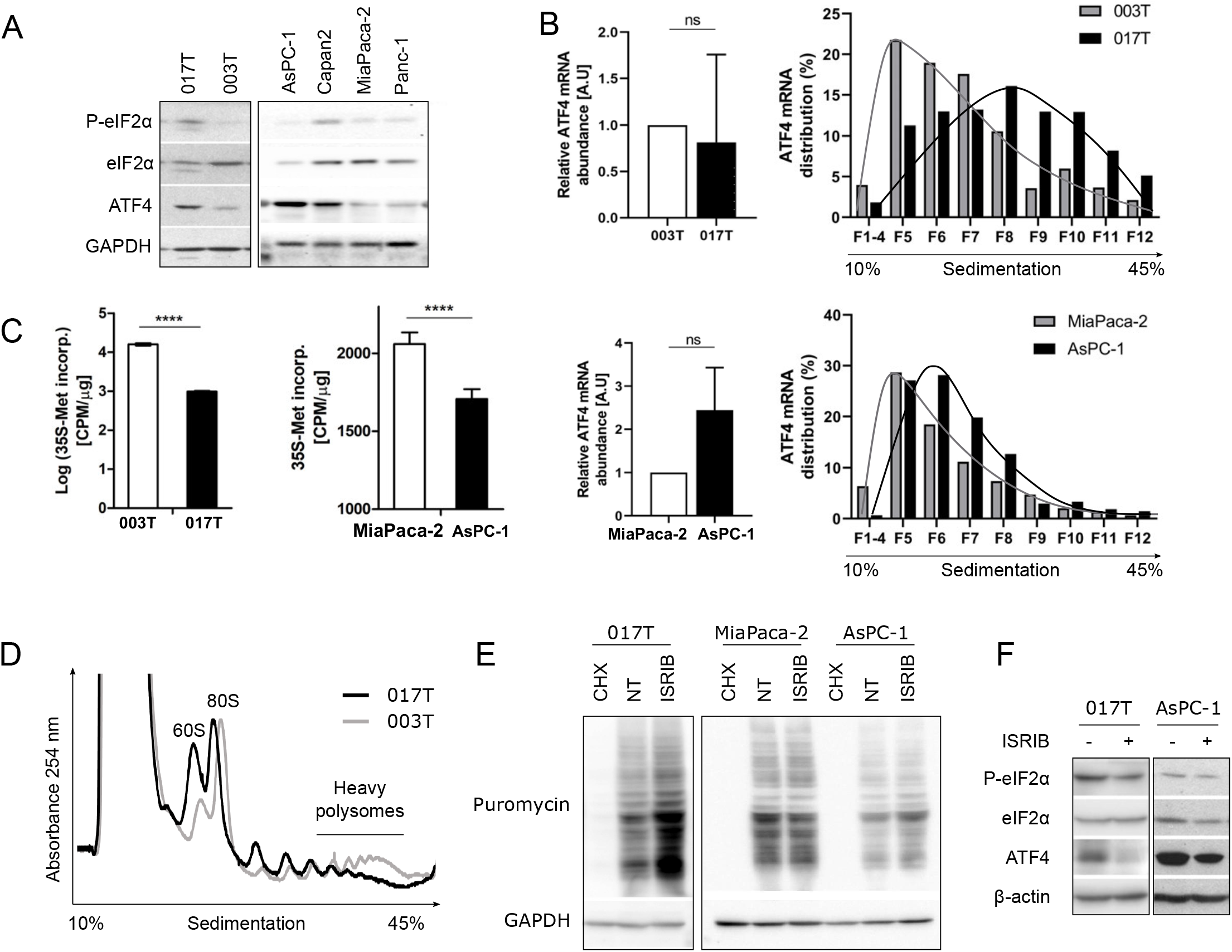
Translation properties of ISR-activated cells. **A)** Western blot analysis of indicated proteins in PDX-derived cell lines (left) and in commonly-used PDA cell lines (right). **B)** The total (left) ATF4 mRNA abundance and its distribution along polysome fractions (right) analyzed by RT-qPCR. **C)** Global mRNA translation rate measured through [^35^S]-methionine incorporation in the 4 cell lines. Data are presented as mean ± SD (n = 3). P-values were calculated using Student’s t-test (****P < 0.0001). **D)** Polysome profiles of PDX-derived cells. Absorbance at 254 nm is shown as a function of sedimentation. **E)** Puromycin incorporation in the indicated cell lines treated with 100 μg/mL CHX for 15 min or 1 μM ISRIB for 1 hr is visualized through western blot. **F)** Western blot analysis of ISR-activated cells treated or not with 1 μM ISRIB for 1 hr.

### ISR-activated cells are resistant to chemotherapies and to apoptosis

ISR and phosphorylation of eIF2α are controlled by kinases (PERK, HRI, PKR and GCN2) and phosphatases (GADD34 and CReP). Unexpectedly, the expression of none of these regulatory enzymes were strongly associated with the IC3 component (Supp. Fig. 2C). Specific stimuli of each kinase were applied on MiaPaca-2 and AsPC-1 to identify potential impairment or activation of a single ISR arm (Fig. 3A). Amino acids starvation (no AA), achieved using Hank’s balanced Salt Solution (HBSS), activates GCN2. Poly(I:C), mimicking viral infection, leads to PKR activation. Arsenite is thought to activate HRI and finally, Tunicamycin (Tuni), an inhibitor of glycosylation, activates PERK in the ER (Taniuchi et al. 2016). Each of these treatments induced eIF2α phosphorylation to a similar extent in both MiaPaca-2 and AsPC-1 cells, although we did not observe ATF4 induction upon Poly(I:C) treatment. The most striking difference was a reduced expression of ATF4 in ISR-activated AsPC-1 cells upon Tunicamycin treatment (Fig. 3A). Looking at the comparative kinetic of ISR induction in 003T and 017T cells, Tunicamycin was found to induce PERK upshifting (due to phosphorylation), ATF4 and BiP expressions within the same time period, although ATF4 induction was weaker in ISR-activated cells as well as PARP cleavage, an apoptosis marker (Fig. 3B). Treatment with other PERK inducers such as BrefeldinA or the BiP inhibitor HA15 (Cerezo et al. 2016) also led to a reduced induction of ATF4 in 017T cells, as compared to 003T cells, despite a similar increase of eIF2α phosphorylation, CHOP and UPR activation (visualized by IRE1 shift and XBP1s expression) (Fig. 3C). PERK inhibitor (GSK) did not decrease ATF4 basal expression in 017T cells, indicating that PERK kinase is not activated under standard culture condition. Conversely, ATF4 abundance was reduced in 017T cells upon ISRIB treatment, as reflected by the elevated relative eIF2α phosphorylation (lower expression of total eIF2α with similar phosphorylation) (Fig. 3C). Concordantly, both ISR-activated cells were more resistant to Tunicamycin than translational reference cells (Fig. 3D). In fact, a deeper look at apoptosis confirmed that neither caspase-3, nor PARP, were cleaved in ISR-activated cells upon sustained ISR induction with HA15, Tunicamycin (Fig. 3E), TRAIL treatment or amino acid starvation (Fig. 3F). Apoptosis resistance of 017T cells upon MG132 treatment, a proteasome inhibitor, was also confirmed by Annexin V labelling (Fig. 3G). Facing this marked resistance to apoptosis, chemotherapeutic agents were tested. Gemcitabine (Gem) in combination with Abraxane (*nab*-Paclitaxel, Abx) is one standard first-line chemotherapy for ECOG0-1 PDA patients (Von Hoff et al. 2013). Gem was previously reported to induce ISR (Palam et al. 2015), but, to our knowledge, ISR stimulation upon Gem/Abx treatment has never been characterized. In MiaPaca-2 cells, Abx induced a much stronger ISR compared to Gem, as evidenced by increased PERK/eIF2α phosphorylation and ATF4 expression (Fig. 3H). Surprisingly, Gem/Abx combination induced ISR similarly to Abx alone, although ATF4 expression was slightly reduced. Comparing AsPC-1 (ISR-activated cells) to MiaPaca-2 cells, Abx and Gem/Abx induced similar DNA damage, as visualized by phospho-γH2AX signals, yet, these treatments induced apoptosis only in MiaPaca-2 cells (Fig. 3H). Finally, chemoresistance of ISR-activated cells was confirmed by cell viability assay showing that Gem and Abx were more cytotoxic on translational reference cells, MiaPaca-2 and 003T, than on ISR-activated cells (Fig. 3I). Taken together, these data highlight an impressive resistance to apoptosis and to chemotherapy of ISR-activated cells.

**Figure 3:**
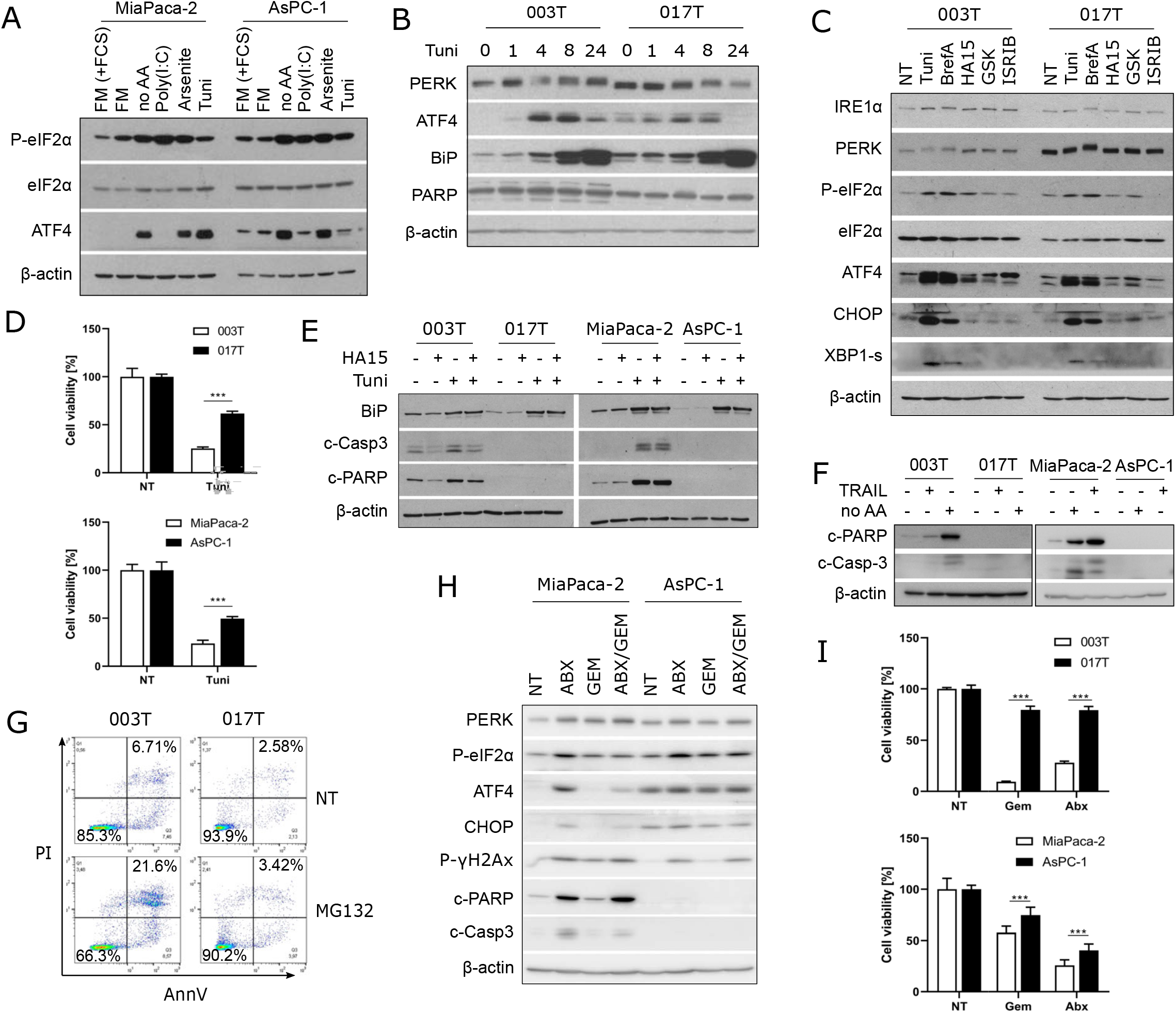
ISR-activated cells are resistant to chemotherapies and apoptosis. **A)** Western blot analysis of indicated proteins. Cells were starved in full media without serum (FM) or without serum nor amino acids (no AA) using HBSS, or transfected with 10 μg/mL Poly(I:C), treated with 500 μM Arsenite or 8 μg/mL Tunicamycin (Tuni) in full media with serum for 6 hrs. **B-C)** Western blot analysis of PDX-derived cell lines treated with **B)** 8 μg/mL Tunicamycin for the indicated time or **C)** 8 μg/mL Tunicamycin, 0.5 μg/mL BrefeldinA (BrefA), 10 μM HA15, 50 nM GSK2606414 (GSK) or 500 nM ISRIB for 4 hrs. **D)** MTT assay with cells treated or not with 1 μg/mL Tunicamycin for 48 hrs in full media. Data are presented as mean ± SD (n = 3). P-values were calculated using 2-way ANOVA (***P < 0.001). **E-F)** Western blot analysis of cells treated **E)** with 1 μM HA15 or 4 μg/mL Tunicamycin, or both for 48 hrs, or **F)** 100 ng/mL TRAIL in full media or starved in HBSS (no AA) for 16 hrs. **G)** Flow cytometry analysis of PDX-derived cells stained with Propidium Iodide (PI) and AnnexinV (AnnV) after treatment with 250 μM MG132 for 6 hrs in full media. **H)** Western blot analysis of MiaPaca-2 and AsPC-1 cells treated with 30 μM Abraxane (ABX) or 1 μM Gemcitabine (GEM), or both, for 24 hrs in full media. **I)** MTT assay with cells treated or not with 100 μM Gemcitabine or 5 μM Abraxane for 48 hrs in full media. Data are presented as mean ± SD (n = 3). P-values were calculated using Student’s t-test (***P < 0.0005).

### ISR-activated cells present low autophagy

Pancreatic cancer requires autophagy for tumor growth (Yang et al. 2018). Interestingly, ATF4 has been demonstrated to transcriptionally induce autophagosome components such as LC3B, Beclin1, ATG12 and ATG3 (B’chir et al. 2013). Thus, autophagy dependence was questioned in ISR-activated cells, and autophagosome abundance was first quantified by LC3B immunofluorescence. Despite the high expression of ATF4, surprisingly, ISR-activated cells showed much lower basal abundance of LC3B puncta, as compared to translational reference cells (Fig. 4A). Upon autophagy inhibition using Chloroquine (CQ), the quantified number of autophagosomes dramatically increased in both cell types, but still remained lower in ISR-activated cells (Fig. 4A, lower panel), again suggesting a lower autophagy in the latter. This was also confirmed through western blot (Fig. 4B), as evidenced by the lower abundance of LC3BII form in ISR-activated cells, compared to translational reference cells. Their autophagic fluxes were also evaluated by quantifying the relative accumulation of LC3BII upon CQ treatment. LC3BII signals of long exposure in ISR-activated cells were compared to those of short exposure in translational reference cells, in order to obtain similar initial signals (see quantification Supp. Fig. 3A). The similar autophagic flux observed in both groups was indicative of a functional autophagy in ISR-activated cells. Combination of amino acid starvation (no AA) with CQ is thought to favor apoptosis due to the inability of cells to generate missing amino acids through autophagy. As we previously observed, ISR-activated cells were resistant to apoptosis induced by amino acids depletion (no AA) and/or CQ (Fig. 4C), which was further confirmed by a cell viability assay (Fig. 4D). Since ISR-activated cells display a high basal expression of ATF4 associated with limited amount of eIF2α, we asked if overexpression of ATF4 was sufficient to induce ISR-activated phenotype in MiaPaca-2 cells, or if increasing the amount of eIF2α would reverse the ISR-activated phenotype (Supp. Fig. 3B-F). Neither eIF2α-overexpressing AsPC-1 cells, which demonstrated enhanced protein synthesis, nor ATF4-overexpressing MiaPaca-2 cells, developed modifications of autophagic capacities, resistance to apoptosis or sensitivity to chemotherapies, as compared to cells transduced with control lenti-vector or parental cells. Together, these data demonstrate that, ISR-activated cells present low autophagic capacities, and that modulation of mRNA translation or expression of ATF4 cannot trigger the ISR-activated cells phenotype, suggesting that ATF4 is rather a consequence of more profound cellular modifications.

**Figure 4:**
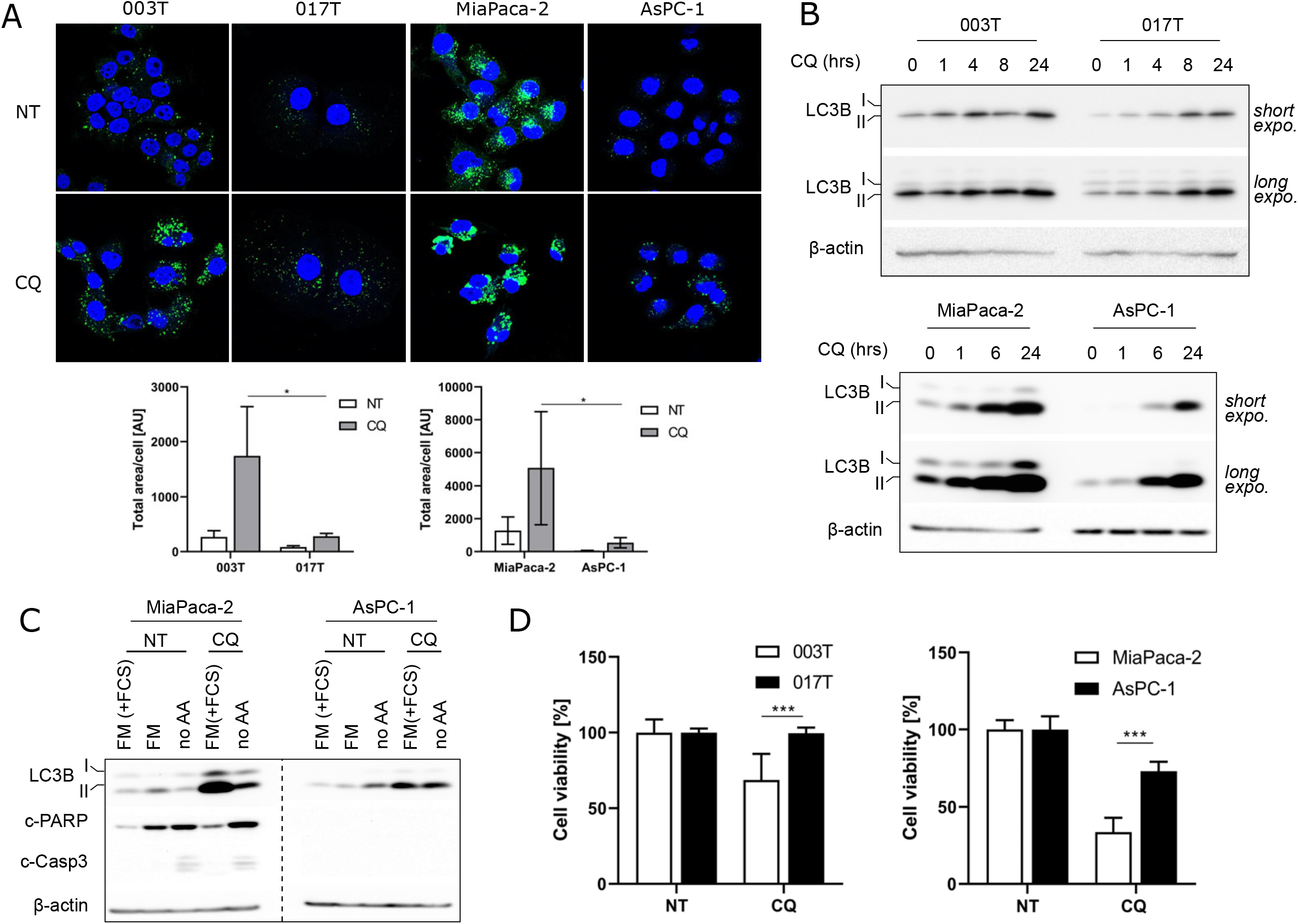
ISR-activated cells have low autophagy. **A)** Immunofluorescence of LC3 in cells treated or not with 50 μM Chloroquine (CQ) for 4 hrs in full media. %Area of LC3 per cell was quantified using ImageJ. **B)** Western blot analysis of indicated proteins in cells treated with 50 μM CQ for the indicated time in full media. Long and short exposures are depicted. **C)** Western blot analysis of cells starved in full media with (FM+FCS) or without serum (FM) or media without amino acids (no AA), or treated with 50 μM CQ in FM or no AA for 24 hrs. **D)** MTT assay with cells treated or not with 50 μM CQfor 48 hrs in full media. Data are presented as mean ± SD (n = 3). P-values were calculated using 2-way ANOVA (***P < 0.001).

### ISR-activated cells show impaired serine biosynthesis

Since ISR-activated cells harbor low autophagic capacities, we thought that the access to exogenous nutrients would be critical for their growth. Thus, cell density was measured 48 hours after incubation in minimal essential media (MEM) containing only essential amino acids (EAA, Fig. 5A). As expected, ISR-activated cells showed lower growth capacity as compared to translational reference cells. Upon amino acid deprivation (no AA), ATF4 induction was observed in all PDA cells, although moderated in 003T cells (Fig. 5B). The sole supplementation for 4 hours with essential amino acids (EAA) was sufficient to reduce ATF4 expression in translational reference cells. Conversely, both essential and non-essential AA (EAA+NEAA) were necessary to decrease ATF4 expression in ISR-activated cells. HBSS supplemented with essential AA and individual non-essential AA allowed the identification of serine as a critical AA for ISR-activated cells (Fig. 5C). Surprisingly, despite the fact that glycine and serine are able to interconvert through the enzymes SHMT1/2 (Yang and Vousden 2016), glycine was not able to reduce ATF4 in ISR-activated cells. This result was confirmed by using full media only lacking serine and glycine (Supp. Fig. 4A). We found that, similarly to ISR-activated cells, Capan-2 cells which had high expression of ATF4 and robust eIF2α phosphorylation in normal culture condition (Fig. 2A), also presented serine dependency (Supp. Fig. 4B-C). Analysis of protein synthesis under nutrient restriction was inversely correlated with ATF4 expression (Fig. 5D). Depletion of NEAA induced marked decrease of puromycin incorporation in ISR-activated AsPC-1, which was fully recovered by the addition of NEAA (serine, proline, glutamate, glycine, asparagine, aspartate, alanine), or serine alone. These data overlapped with cell growth, where the single addition of serine to media containing only EAA could restore AsPC-1 cell proliferation (Fig. 5E). MiaPaca-2 cells showed little changes of protein synthesis and viability under similar growth conditions. These results suggest an impairment of serine biosynthesis pathway (or SBP) in ISR-activated cells (Fig. 5F). Thus, we quantified using RT-qPCR mRNAs encoding the phosphoglycerate dehydrogenase (PHGDH), the phosphoserine aminotransferase (PSAT1) and, to a lesser extent, the phosphoserine phosphatase (PSPH), and found that these mRNAs are reduced in ISR-activated cells, as compared to translational reference cells (Fig. 5G). Furthermore, treatment with a PHGDH inhibitor (CBR-5884) in media lacking serine (MEM) induced ATF4 expression in translational reference cells, whereas it had no effect in ISR-activated cells (Fig. 5H). Finally, PHGDH depletion in ISR-activated cells was validated at the protein level (Fig. 5I). As in the presence of EAA, glycine had no effect on ATF4 expression in ISR-activated cells, we thought that fueling the folate cycle with formate could sustain serine-dependent cell proliferation (Labuschagne et al. 2014). Indeed, addition of formate and glycine to MEM (EAA) favors the reduction of ATF4 similar to serine in AsPC-1. Furthermore, it partially restored cell growth whereas glycine alone had no effect (Supp. Fig. 4D-E). Impairment of SBP appears counterintuitive considering the role of ATF4 as transcriptional activator of this pathway (Ye et al. 2012). Thus, the inability of ATF4 to stimulate the expression of PHGDH was questioned in ISR-activated cells. As demethylating agent 5-azacytidine was previously shown, in AsPC-1 cells, to favor expression of specific genes, including MMPs (Sato et al. 2003), we applied a similar strategy and observed a consistent re-expression of PHGDH, which was further induced upon EAA removal (Fig. 5J). Finally, we validated by immunohistochemistry (IHC) on the initial PDX 003T and 0017T, as well as on orthotopically-grafted MiaPaca-2 and AsPC-1 cells into nude mice, a lower expression of PHGDH in ISR-activated tumors, as compared to translational reference tumors. Interestingly, in ISR-activated tumors, stromal expression of PHGDH appeared to be increased, especially in fibroblastic cells (Supp. Fig. 5). Altogether, these data demonstrate that ISR-activated cells display serine auxotrophy due to the lack of PHGDH expression. Moreover, the single serine supplementation is sufficient to rescue protein synthesis and cell growth, while other NEAA are not required.

**Figure 5:**
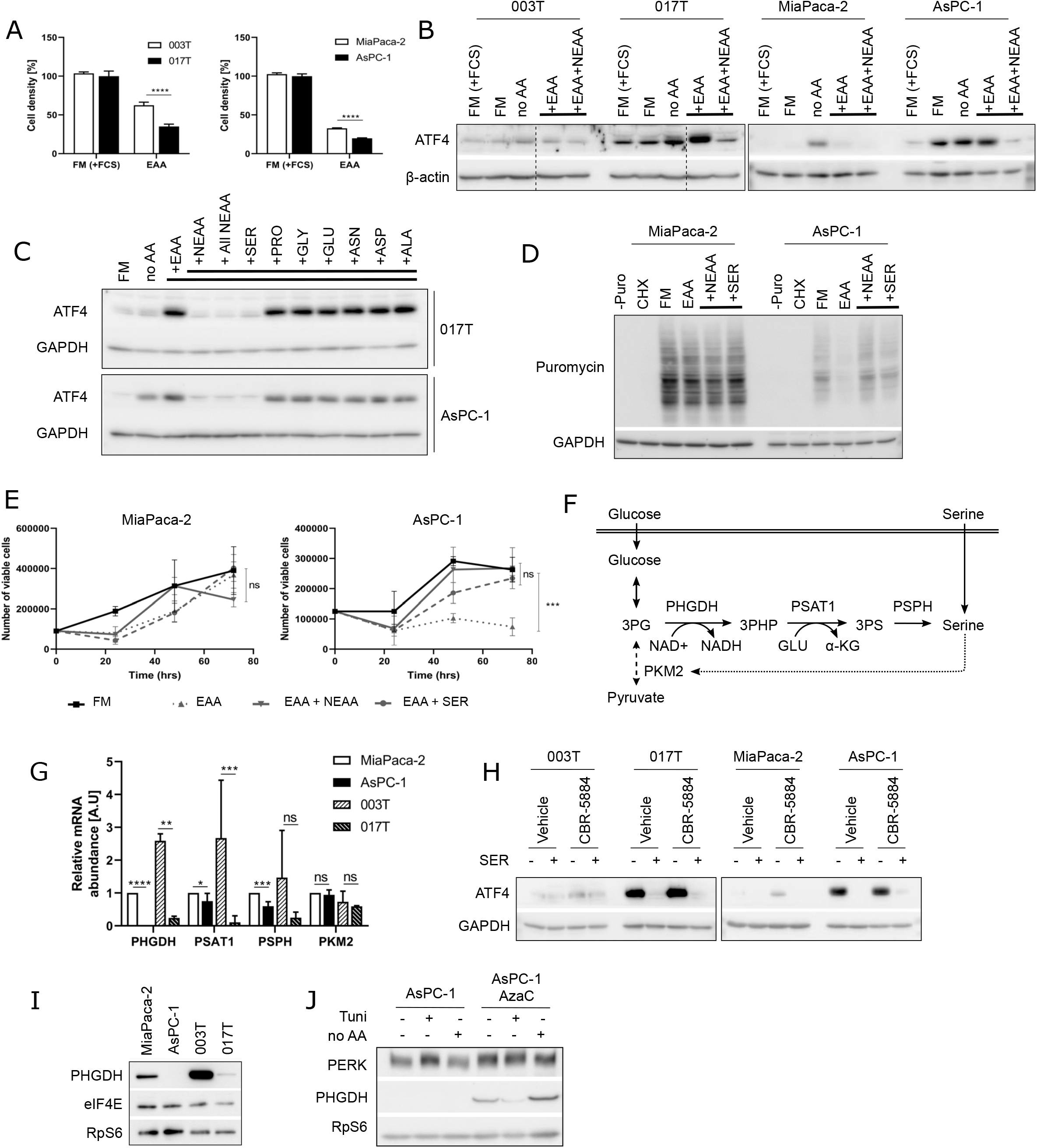
ISR-activated cells have impaired serine biosynthesis pathway. **A)** Crystal Violet assay with cells in full media with serum (FM+FCS) or in media only with EAA for 48 hrs. Data are presented as mean ± SD (n = 3). P-values were calculated using 2-way ANOVA (****P < 0.0001). **B-C)** Western blot analysis of indicated proteins. Cells were grown in FM with serum or starved in media without amino acids (no AA) for 2 hrs then **B)** essential amino acids (EAA) and non-essential amino acids (NEAA) were added to the media for 4 additional hours or **C)** 100 μM of the indicated non-essential amino acids (serine, SER; proline, PRO; glycine, GLY; glutamate, GLU; asparagine, ASN; aspartate, ASP; alanine, ALA) were added with EAA to HBSS for 4 additional hours. As a control, all individual NEAA were added simultaneously to the media (All NEAA). **D)** Puromycin incorporation in cells treated with 100 μg/mL CHX for 15 min in full media, starved in media with only EAA, supplemented with NEAA or serine (SER) for 1 hr visualized through western blot. **E)** Proliferation assay of MiaPaca-2 and AsPC-1 cells cultured in the indicated medium. Data are presented as mean ± SD (n = 3). P-values were calculated using 2-way ANOVA (***P < 0.001). **F)** Schematic of the serine biosynthesis pathway. Phosphoglycerate deshydrogenase (PHGDH), Phosphoserine aminotransferase (PSAT1), Phosphoserine phosphatase (PSPH) are 3 enzymes implicated in serine synthesis from glucose. Pyruvate kinase (PKM2) catalyzes the last step of pyruvate synthesis from glucose. High concentration of serine is able to positively regulate PKM2. **G)** mRNA abundance of the indicated genes was analyzed through RTqPCR. Data are presented as mean ± SD (n = 3). P-values were calculated using 2-way ANOVA (*P< 0.05, **P<0.005, ***P < 0.001, ****P<0.0001). **H-J)** Western blot analysis of **H)** cells treated with 30 μM CBR-5884 (PHGDH inhibitor) for 6 hrs in media without NEAA (MEM) supplemented or not with 100 μM serine (SER). **I)** All four cell lines in full media with serum **J)** AsPC-1 cells cultured with 1uM Azacitidine (AzaC) for 3 serial passages or not. Then, treated with 8 μg/mL Tunicamycin (Tuni) or starved in media without amino acids (no AA) for 24h.

### Rewiring of the transsulfuration pathway limits serine consumption in ISR-activated cells

Serine is a key metabolite for the transsulfuration (TSS) pathway, one essential metabolic pathway for glutathione formation. Abundance of reduced glutathione is critical to maintain low cellular concentration of reactive oxygen species (ROS) and to disable oxidation of lipids, proteins and nucleic acids in PDA (DeNicola et al. 2011). The TSS pathway starts by the conversion of homocysteine to cystathionine through serine incorporation by the cystathionine β synthase (CBS, Fig. 6A). Considering the serine dependency of ISR-activated cells, expression of enzymes from the TSS pathway was examined by RT-qPCR (Fig. 6B). All mRNA, protein and activity of CBS were drastically down-regulated in ISR-activated cells, as compared to translational reference cells (Fig. 6B-D). Interestingly, and despite the absence of CBS, we found that AsPC-1 cells had a lower basal ROS level than MiaPaca-2 cells. Moreover, ROS level increased in MiaPaCa-2, but not in AsPC-1, cells when cultured in MEM (EAA) (Fig. 6E). In addition, culturing AsPC-1 cells with NAC, a potent ROS scavenger, had no effect on cell growth (Supp. Fig. 4F). Thus, TSS pathway appears functional in ISR-activated cells without supplementation in serine. As a consequence, we hypothesized that extracellular cysteine is needed to bypass CBS depletion in order to generate glutathione in these cells (Fig. 6A). In fact, ATF4 expression was induced after cysteine depletion in the growth media of ISR-activated cells, whereas translational reference cells, bearing an active CBS, showed no induction of ATF4 (Fig. 6F). An inhibitor of cysteine transporter xCT, Erastin, had similar effect by limiting the cellular import of cysteine (Fig. 6G). These data overlapped with cell growth, where the cysteine depleted media or Erastin treatment inhibited proliferation of ISR-activated AsPC-1 cells (Fig. 6H). Finally, using IHC, we showed a lower CBS protein expression in 017T PDX and AsPC-1-grafted tumors, as compared to translational reference tumors, validating our *in vitro* data (Supp. Fig. 5). All together, these data show that ISR-activated cells present an impairment of the first step of TSS pathway requiring serine, thereby revealing their cysteine dependency.

**Figure 6:**
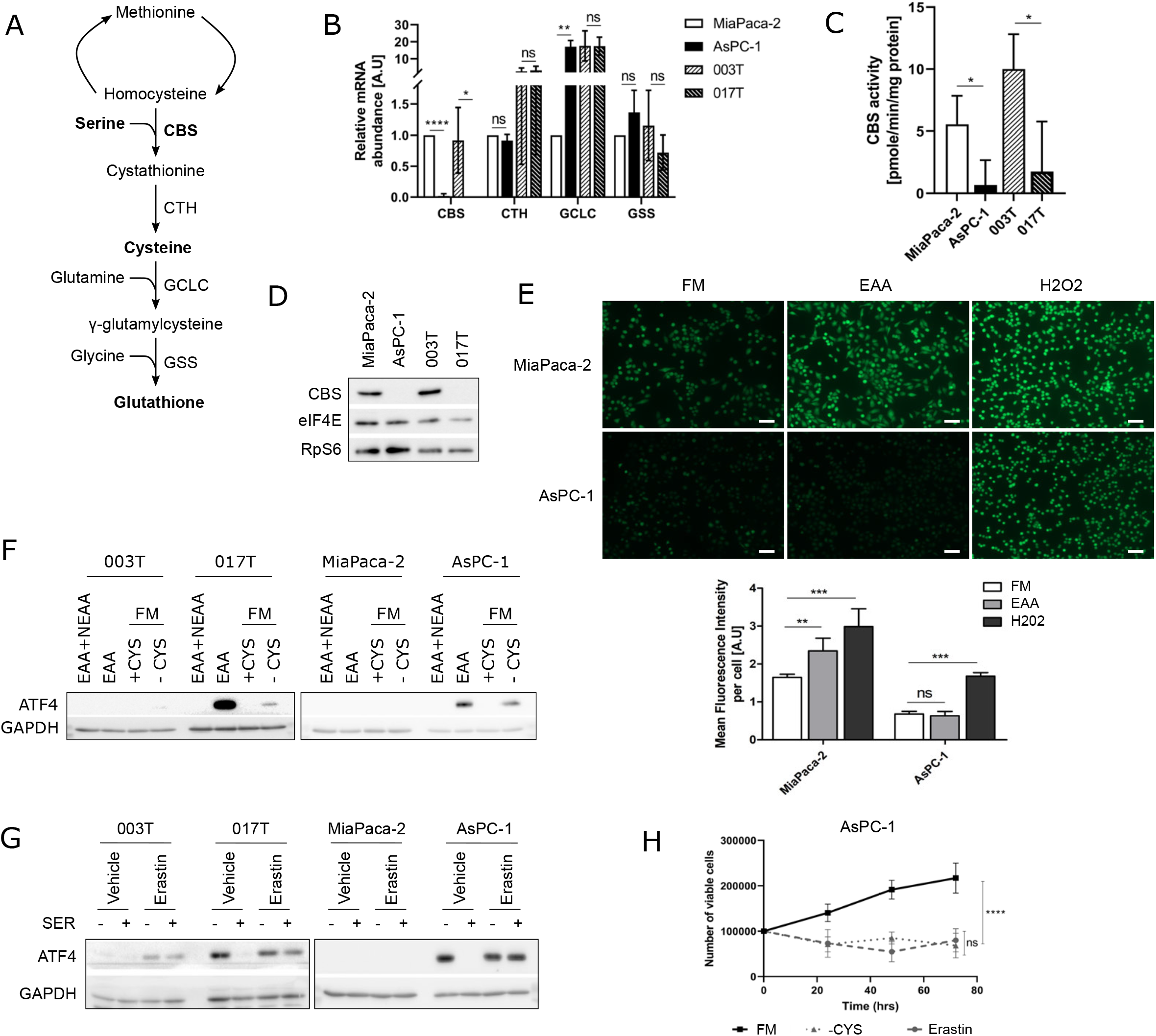
Rewiring of transsulfuration pathway limits serine consumption in ISR-activated cells. **A)** Schematic of the transsulfuration pathway. Cystathionine-β-synthase (CBS), cystathionine-γ-lyase (CTH), Glutamate-Cysteine Ligase (GCLC) and Glutathione synthase (GSS) are enzymes implicated in glutathione synthesis from Homocysteine. **B)** mRNA abundance of the indicated genes was analyzed through RTqPCR. Data are presented as mean ± SD (n = 3). P-values were calculated using unpaired t-test (*P< 0.05, **P<0.005, ****P<0.0001). **C)** CBS activity measured in all four cell lines, and normalized with the protein quantity. Data are presented as mean ± SD (n = 3). P-values were calculated using paired t-test (*P < 0.01). **D)** Western blot analysis of all four cell lines in full media with serum **E)** DCFDA staining of cells in the indicated condition (up). The scale bars correspond to 100 μm. Mean fluorescence intensity quantification of the DCFDA staining using the ZEN software (down). P-values were calculated using 2-way ANOVA (**P< 0.005; ***P< 0.001). **F-G)** Western blot analysis of cells **F)** incubated in MEM media containing only EAA, or supplemented with NEAA (EAA+NEAA); Full Media FM without (-CYS) or with 50 μM cysteine for 6 hrs (+CYS); **G)** treated with 10 μM Erastin (xCT inhibitor) in media with only EAA (MEM) supplemented or not with 100 μM serine for 6 hrs. **H)** Proliferation assay of AsPC-1 cells cultured in the indicated medium: full media (FM), media without cysteine (- CYS) and FM with 2 μM Erastin (Erastin). Data are presented as mean ± SD (n = 3). P-values were calculated using 2-way ANOVA (Turkey’s multiple comparison test) (****P < 0.0001).

## DISCUSSION

Alteration of protein synthesis in cancer cells is now a well admitted notion, further supported by the activation of many signaling pathways. In addition, many translation initiation factors favor cell transformation, whereas some translational regulators, such as PDCD4 and 4E-BP1, are considered as tumor suppressors (Bhat et al. 2015; Martineau et al. 2013). Despite all these evidences, studying cancer cell mRNA translation *in vivo*, in the context of the whole tumor environment, has been extremely challenging. Until recently, many technical limitations have blocked the way to solve this unanswered question (King and Gerber 2014). Here, we succeeded in performing a genome-wide analysis of translated mRNA to identify modulation of protein synthesis in PDA solid tumors, using PDX samples. These PDA avatars gather the advantages of maintaining human cancer cell heterogeneity and tumor histology, while human stroma is replaced by a similar mouse stroma. Using unbiased bioinformatics approaches, translatome analysis uncovered a subset of tumors, distinct from transcriptome-based basal-like and classical subtypes. These tumors are characterized by a low protein synthesis rate and a persistent activation of the ISR pathway, as evidenced by translational activation of mRNA encoding ATF4, ATF5 and JUN. Functional characterization of the ISR-activated PDX-derived cancer cells revealed unforeseen features. At first, ISR-activated cells are refractory to most chemotherapies used on PDA patients and display a complete resistance to apoptosis. Then, ISR-activated cells fail to activate specific ATF4-dependent transcriptional programs, such as autophagy, apoptosis and SBP. Finally, ISR-activated cells show a dual amino acid auxotrophy to serine and cysteine. The defect in SBP, arising from the lack of PHGDH, coincides with a rewiring of TSS pathway, where serine is no longer used to produce cysteine, due to the absence of CBS. Consequently, growth of ISR-activated cells entirely relies on external sources of serine and cysteine.

Identification of ISR-activated tumors echo with a recent publication, where authors discovered a serine dependency of certain PDA cells, by examining the metabolic support provided by neurons in tumors (Banh et al. 2020). Our study is based on a reversed and unbiased approach, whereby translatome analysis allowed the identification of one original tumor subtype. This common identification further confirms the existence of such particular subtype of PDA cancer cells, for which we had further characterized above-mentioned critical features (i.e., cysteine-dependency, low autophagy, and drug-resistance). Moreover, the translatome-based method might be applicable to future molecular profiling of tumors.

Defining pancreatic cancer vulnerabilities is the holy grail of molecular profiling in order to improve patient treatment, objective response rate and overall survival (Singh and O’Reilly 2020). The best characterized molecular profiling arised from exome sequencing and the identification of germline mutation of BRCA1/2 in 14% of PDA patients. BRCAness patients have defective DNA repair and show high vulnerability to platinum-based chemotherapy (Golan et al. 2017). Subsequent clinical trials have established Olaparib as a viable option for maintenance therapy of BRCAness metastatic patients (Golan et al. 2019). Here, translatome analysis identified ISR-activated cancer cells. ISR-activated phenotype is associated with resistance to apoptosis after treatment with Gemcitabine, *nab*-Paclitaxel, TRAIL. The serine auxotrophy is likely the cause of the ISR-activated phenotype, as PHGDH inhibition led to permanent ISR activation. On the contrary, targeting eIF2α phosphorylation using ISRIB or overexpressing eIF2α can restore protein synthesis, but has little impact on the ISR-activated phenotype. This indicates that the permanent activation of ISR is likely a consequence of serine auxotrophy.

ISR-activated cells are also resistant to the inhibitor of autophagosome formation, Chloroquine. Strikingly, many reports identified autophagy as an essential driver for pancreatic cancer development in mouse models including PDX (Yang et al. 2018), despite poor results on patient survival of chloroquine in clinical trials (NCT01273805). Previously, ATF4 expression has been characterized as a transcriptional activator for many autophagosome components including LC3B, Beclin1, ATG12 and ATG3 (B’chir et al. 2013). Surprisingly, ISR-activated cells had low autophagic capacity despite a permanent ATF4 expression. Overall, ISR-activated cancer cells fail to activate some ATF4-dependent transcriptional programs, including SBP enzymes (Ye et al. 2012), apoptotic pathway (Pakos-Zebrucka et al. 2016) and autophagy (as observed in Figs. 3–5). The ATF4-coordinated stress response is thought to allow environmental adaptation by enhancing appropriate cell processes. For example, under limited supplies of amino acids, the GNC2-ATF4 axis favors both autophagy and amino acid synthesis in order to respond to cellular demands. In ISR-activated cells, the impairment of ATF4-coordinated stress response is likely to induce their dual auxotrophy phenotype, and to result in the persistent eIF2 phosphorylation and subsequent decrease of global protein synthesis (see Fig 2).

Pancreatic cancer cells are extremely sensitive to extracellular nutrient availability, especially non-essential amino acids (NEAA). Dependencies on glutamine (Son et al. 2013), alanine (Sousa et al. 2016), proline (Olivares et al. 2017), cysteine (Badgley et al. 2020) and asparagine (Dufour et al. 2012), have been reported and highlight the importance of metabolic reprogramming of PDA cancer cells. This fact contrasts with the hypoxic and nutrient deprived microenvironment of PDA. Cancer-associated fibroblasts (CAF) can support PDA survival by providing some components, such as alanine (Sousa et al. 2016) or collagen as a source of proline (Olivares et al. 2017). Despite important resistance of ISR-activated cells, serine auxotrophy emerges as a weakness, indicating the existence of another NEAA dependency for PDA. Serine synthesis has not been observed in CAF, which remain the most abundant stromal cells in PDA (Sousa et al. 2016; Banh et al. 2020). Nonetheless, we observed an enhanced PHGDH expression in fibroblastic, αSMA-expressing, stromal cells from ISR-activated PDX and grafted AsPC-1, as compared to other tumors (Supp. Fig. 5). This suggests a potential modification of CAF metabolic capacity in such conditions. Among stromal cells, adipocytes could represent a potential source of NEAA including serine (Green et al. 2016). The abundance of adipocytes in the pancreas increases in elderly, especially in the context of obesity and diabetes, which are risk factors for PDA (Molina-Montes et al. 2020). M2 macrophages are the most abundant immune-related stromal cells found in PDA tumors. M2 macrophages reshape anti-tumor immune response and enhance fibrosis (Zhu et al. 2017). PHGDH, the enzyme catalyzing the first step of SBP has recently been identified as a metabolic marker for M2 macrophage polarization (Wilson et al. 2020). Therefore, it is tempting to speculate that M2 could further support tumor progression by providing essential metabolites. Finally, neurons have also been recently reported to produce serine. Through NGF secretion, PDA cancer cells modify axons guidance, allowing intratumoral release of serine (Banh et al. 2020). Development of PHGDH-negative or serine-auxotrophic tumors is favored in certain serine-rich tissues (pancreas or brain) but appears more restricted in mammary gland (Sullivan et al. 2019). Understanding how these cancer cells modify their microenvironment as a serine provider should guide future investigations. As all the above-mentioned cells could participate to this process, especially fibroblastic cells expressing PHGDH, many routes are likely to emerge.

Pancreatic cancer cells dependency on NEAA can constitute a druggable weakness. For example, asparagine restriction induced using a recombinant asparaginase treatment led to reduced PDA tumor growth, although this treatment also induced a translational reprogramming and survival of tumor cells, in part through activation of the GNC2-ATF4 axis (Pathria et al. 2019). Combining asparaginase to GCN2 inhibitor showed synergistic anti-proliferative effect by reducing ATF4 and favoring apoptosis (Nakamura et al. 2018).. Interestingly, a recent Phase IIb clinical trial using Gemcitabine or mFOLFOX6 in combination with erythrocyte-encapsulated asparaginase has shown very encouraging results by improving overall and progression free survival of PDA patients in second-line treatment (Hammel et al. 2020). Altogether, these data highlight the potential of identifying PDA amino-acid dependency as metabolic weakness, whose targeting may open therapeutic windows for precision medicine.

## EXPERIMENTAL PROCEDURES

### PDX polysomes purification

Fragments of 27 pancreatic PDX (Nicolle et al. 2017) were grounded in liquid nitrogen using a BioPulverizer. Around 100g of tumor powder were lysed in 350 μL hypotonic lysis buffer (5 mM Tris-HCl pH 7.5, 2.5 mM MgCl2, 1.5 mM KCl, 100 μg/ml cycloheximide, 2 mM DTT, 0.5% Triton X-100, 0.5% sodium deoxycholate and 1mM Ribonucleoside vanadyl complexes). 250 μL of lysates were loaded onto a 5 mL discontinuous 5%-34%-55% sucrose gradient (20 mM HEPES-KOH pH 7.6, 100 mM KCl, 5 mM MgCl2) and centrifuged at 45,000 rpm (SW 55 Ti rotor, Beckman Coulter, Inc.) (adapted from Liang et al. 2018) for 45 minutes at 4°C per groups of 6 including a cellular control for each batch. The absorbance at 254 nm was continuously recorded using an ISCO fractionator (Teledyne ISCO) during fractionation. Fraction 8 and 9 containing mRNAs associated to more than 3 ribosomes, were pooled and purified using TRIzol-LS (Thermo Fisher Scientific) in parallel to 50 μL of the total lysate. The quality (DV200> 60%) and the quantity (concentration) of all mRNA samples were evaluated using a Fragment Analyser (Agilent).

### RNA sequencing

Libraries were prepared with NEBNext® Ultra™ II Directional RNA Library Prep Kit for Illumina according to the supplier recommendations (NEB). Briefly, the key stages of this protocol were successively, the removal of ribosomal RNA fraction from 80 ng of total RNA using the RiboCop rRNA Depletion Kit V1.2 (Human/Mouse/Rat) from Lexogen, a fragmentation using divalent cations under elevated temperature to obtain approximately 300bp fragments, double strand cDNA synthesis, using reverse transcriptase and random primers, and finally Illumina adapters ligation and cDNA library amplification by PCR for sequencing. Sequencing was then carried out in paired-end 75b mode with an Ilumina HiSeq 4000. Base calling was performed using the Real-Time Analysis software sequence pipeline (version 2.7.7) from Illumina with default parameters.

### Translatome and Independent Component Analysis

Gene counts were normalized using TMM. The residuals of the linear model between the polysome-RNA-counts and total-RNA-counts (corresponding to Translation efficiency) were computed and subsequent analyses were only performed on the residuals of a subset of genes for which relevant linear models were obtained. This was done by selecting models with: constant error variance (using the Score Test with a minimum p-value of 1%), normally distributed residuals (using Shapiro-Wilk’s test with a minimum p-value of 1%) and the absence of outliers (using Bonferroni outlier test with alpha=5%). ICA was performed on the 50% most variable (using inter quartile range) selected genes residuals using the JADE algorithm.

### Cell culture

AsPC-1, Panc-1, and Capan-2 cells were obtained from American Type Culture Collection (ATCC) and cultured in RPMI (#R0883,MilliporeSigma), and MiaPaca-2 in high-glucose DMEM (#D6429, MilliporeSigma), supplemented with 10% (v/v) fetal calf serum (ThermoFisher Scientific), 2mM L-Glutamine (GLN) and 1% (v/v) penicillin/streptomycin (Sigma-Aldrich) at 37°C and 5% CO2. Cells derived from pancreatic PDX (003T and 017T) were grown in Serum Free Ductal Media (SFDM) as described (Nicolle et al. 2017).

500,000 cells were seeded in 6-well-plates and grown overnight. Cells were treated or not with indicated concentration of drugs in full media, HBSS (containing no AA, #H6648, MilliporeSigma), MEM (containing no NEAA, #M2279, MilliporeSigma) or DMEM (cysteine-, methionine-, glutamine-free media #D0422, MilliporeSigma) supplemented or not with essential/non-essential amino acid mixtures (#M5550, #M7145, MilliporeSigma), or individual amino acids. L-alanine, L-aspartate, L-asparagine, L-glutamate, glycine, L-proline, L-serine, L-methionine and L-cystine were purchased from Sigma-Aldrich.

### Cell polysome profiling and RNA isolation

Cells were cultivated in 15 mm culture dishes. At ~80% confluence, cells were treated with 100 μM cycloheximide (CHX) for 5 minutes before being harvested on ice. Cells were washed twice with cold PBS with 100 μM cycloheximide and lysed in the hypotonic lysis buffer described above. After normalizing the RNA concentration, lysates were loaded onto a 5 mL linear 10%-45% gradient, and centrifuged at 45,000 rpm for 45 minutes at 4°C. The absorbance at 254 nm was continuously recorded using an ISCO fractionator and 12 fractions of ~400 μL were collected along the fractionation. The first four fractions were pooled before mRNA isolation of all fractions and the total lysates, using TRIzol-LS (Thermo Fisher Scientific).

### Compound used

Chemical compounds used in cell culture experiments are listed below: ISRIB (#SML-0843), Poly(I:C) (#I3036), Tunicamycin (#T7765), MG132 (#C2211), BrefeldinA (#B7651) and GSK2606414 (#516535), Chloroquine (#C6628), CBR-5884 (SML-1656) and Erastin (#329600) were obtained from Sigma Aldrich and Trail (#310-04) from Peprotech. BiP inhibitor HA15 was described previously (Cerezo et al. 2016). Compounds were used in full media with serum, unless indicated. Nab-paclitaxel/ABRAXANE (Celgene), Gemcitabine (Sandoz) and Azacitidine (Vidaza, Celgene) were provided by the Pharmacy of “Institut Universitaire du Cancer Toulouse”.

### RTqPCR

Concentrations of RNAs were measured using a NanoDrop (ND-1000 from Thermo Fisher Scienrific). 1 μg of RNA was first reverse transcribed with RevertAid H Minus Reverse Transcriptase (Invitrogen) using random hexamer primers (Invitrogen) according to the manufacturer’s instruction. qPCR was carried out on a StepOne Plus (Applied Biosystems, Life Technologies) using SsoFast EvaGreen Supermix (Biorad) and primer concentration at 0.5 μM. For mRNA distribution across the linear gradient, equal volume of RNAs were used to perform RT-qPCR. The mRNA expression fold-change was normalized using the delta-delta Ct method with RPS16 or GAPDH as a control gene. Primers specific to human mRNAs were purchased from Integrated DNA Technologies (IDT) and are listed on the supplementary table 1.

### Preparation of Cell Extract and Western blot

Cells were washed twice in cold PBS on ice and lysed in 50 mM Tris-HCl pH 7.5, 150 mM NaCl, 2 mM EDTA, 1% NP40, 0.1% SDS and 0.5% sodium deoxycholate supplemented with EDTA-free protease inhibitor and PhosSTOP phosphatase inhibitor (Roche Applied Science), then centrifuged at 13,000 rpm for 5 minutes. Protein concentrations were measured using Protein assay reagent (Biorad) and equal amount of proteins were loaded into a 6%-15% SDS-polyacrylamide gradient gel and transferred onto nitrocellulose membrane (BioTraceNT; Pall Corp.). Membranes were washed in Tris Buffer Saline with 0.1% Tween 20 (TBS-T) and saturated in TBS-T with 5% non-fat dry milk for 20 minutes and incubated overnight in the primary antibody solution (TBS-T with 5% BSA). The following antibodies were used to detect proteins of interest: anti-ATF4 (#11815), anti-eIF2α (#2103), anti-phospho-eIF2α (#3398), anti-CHOP (#2895), anti-cleaved-caspase-3 (#9579S), anti-PARP (#9542S), anti-BiP (#3177S), anti-PERK (#3192S), anti-eIF4E (#2067S) and anti-LC3 (#3868) were purchased from Cell signalling Technology, anti-GAPDH (#25778), anti RpS6 (sc-74459) was purchased from Santa Cruz Biotechnology, and anti-β-actin (clone AC-74), anti PHGDH (HPA021241) were purchased from MilliporeSigma, and anti-CBS (ab140600) was purchased from Abcam. Mouse (#31430) and rabbit (#31460) secondary antibodies from Pierce were used at 1:10000. Signals were revealed by chemoluminescence using ECL (RevelBlot, Ozyme) and captured using Chemidoc imager (Bio-Rad). When indicated, LC3II was quantified using ImageJ (NIH). For each cell line, blots with equivalent LC3II exposure of the untreated condition were quantified.

### SUnSET assay and [^35^S]-Methionine incorporation

500,000 cells were seeded in 6-well plates and grown overnight. For SUnSET assays, cells were incubated with 1 μg/mL Puromycin (Sigma-Aldrich) during 10 minutes before harvesting cells. Protein synthesis rate was visualized by western blot, using the anti-Puromycin (#MABE343; 1/5000) from Merck Millipore. Cycloheximide was added 5 minutes before Puromycin and served as negative control. For [^35^S]-methionine metabolic labeling, cells were incubated for one hour in methionine- and cysteine-free DMEM (#D0422, MilliporeSigma) with 10% dialysed serum (#26400044, Gibco). The medium was then replaced with methionine- and cysteine-free DMEM containing [^35^S]-protein labeling mix (20 μCi/mL, EasyTag EXPRESS35S, Perling Elmer). After 30 minutes, cells were washed with cold PBS and lysed in buffer (see Preparation of Cell Extract and Western Blot), and radioactivity incorporated into the TCA precipitable material was measured. Radioactivity (CPM) was normalized on protein concentration.

### Viability assay

Cell abundance was measured by MTT assay or Crystal violet staining. 6000 cells were plated in 96-well plates, and grown overnight. Cells were treated with indicated drugs in at least quadruplicates. 48 hours later, MTT was added to each wells and incubated at 37°C during 3 hours. After eliminating the media, formazan crystals were dissolved in 100 μL of DMSO. For crystal violet staining, cells were fixed with 3.7% formalin then incubated in crystal violet for 20 minutes. After cell wash with H2O, cells were lysed in 10% acetic acid. The absorbance of each well was read at 570 nm using a microplate spectrophotometer Mithras (Berthold Technology).

### Flow cytometry for apoptosis detection

Cells were plated in 6-well culture plates and were grown overnight. Cells were treated with 250 μM MG132 for 6 hours. Cells were washed twice with cold PBS, and then resuspended in 1x AnnexinV Binding buffer (#556454, BDbioscience) at a concentration of 1.10e6 cells/mL. 5 μL of FITC AnnexinV (#556420, BDbioscience) and 10 μL of 50 μg/mL Propidium Iodide (#556463, BDbioscience) were added to 100 μL of each cell suspension, and were incubated for 15 minutes at RT. 400 μL of 1x AnnexinV Binding buffer was added to each tube and cells were analyzed on a MACSQuant VYB (Miltenyi Biotec). A minimum of 10 000 cells were analyzed per condition. Data analysis were carried out using FlowJo v10.

### Proliferation assay

~100,000 cells were seeded in 12 well-plates in complete media. Once cells adhered to the plate, complete media was gently removed and cells were washed once with PBS to eliminate the serum. Cells were cultivated in the indicated medium in triplicates. After 24, 48 and 72 hours, cells were counted in 0.2% Trypan Blue solution (MilliporeSigma) using Countness II Automated Cell Counter (Applied Biosystems, Life Technologies). Only viable cells were taken in account.

### Immunofluorescence and microscopy

Cells were grown on 15 mm x 15 mm coverslip and treated with 50 μM Chloroquine for 4 hours. After washing twice with cold PBS on ice, cells were fixed with 3.7% formaldehyde for 10 minutes at room temperature (RT), then permeabilized and saturated in PBS supplemented with 3% BSA and 0.01% saponin during 30 minutes at RT. Cells were incubated in anti-LC3 antibody (#PM036 from MBL International Corp.) at dilution 1:800 for 45 minutes, then in secondary antibody conjugated with Alexa Fluor 488 (Invitrogen) at 1:1000 for 30 minutes after cell wash. Cells were incubated in 100 ng/mL DAPI for 10 minutes before mounting slides using the Fluorescent mounting medium (DAKO). Cells were visualized on Zeiss LSM780 confocal microscope using the ZEN software. Images were captured arbitrarily from four different spots using the 63x /1.4 objective, and the %Area of LC3 per cells were quantified using ImageJ (NIH). A color threshold was applied to quantify the total areas of green dots, and were normalized on the number of cells in the field. The mean was calculated for each condition, and the ratio [chloroquine-treated/untreated] to obtain the autophagic flux.

### CBS enzymatic activity

Cells were seeded on a 150 mm diameter tissue culture plate. At ~80% confluency, cells were washed twice with cold PBS and were collected in tubes, centrifuged at 2000 rpm for 10 minutes. Supernatents were eliminated and cell pellets were flashed freeze and conserved at −80°C. CBS activity in cells were measured using the CBS activity fluorometric assay kit from BioVision (#K998) following the manufacturer’s instruction. Fluorescence (Ex/Em 368/460 nm) was measured using Clariostar (BMG Labtech) during 30 minutes.

### DCFDA staining

Cells were seeded on a 6-well plate. At ~80% confluency, cells were put in the indicated media for 2 hours and then incubated with 10 μM DCFDA dye at 37°C without CO2 for 1 hr. Cells were visualized on a videomicroscope Cell Observer using the ZEN software. Images were captured arbitrarily from three different spots using the 10x /10.3 objective, and the Mean Fluorescence Intensity were quantified through ZEN and normalized on the number of cells quantified using Image J (NIH).

### Statistics

Statistical significance was determined using two-way ANOVA or paired/unpaired t-test P-value depending on the experiments. All values are mean ± SD. A P-value <0.05 was considered statistically significant (*P<0.05; **P<0.005; ***P<0.0005; ****P<0.00005).

## ACKNOWLEDGEMENTS

Authors acknowledge Jean-Emmanuel Sarry, Sophie Vasseur, Jean-Charles Portais for helpful discussion on metabolism; Dominique Weil and Pierre Fafournoux for constructive comments on translational control; Sophie Peria from IUCT Pharmacy for providing Gemcitabine, Nab-Paclitaxel and Azacitidine. Loïc Van Den Berghe and Christele Segura for viral particle production (Pôle Technologique CRCT, U1037).

## AUTHOR’S CONTRIBUTION

S.S., R.S., Ch.J., J.R., A.B., J.S. and Y.M. designed and carried experiments. R.N. analyzed RNAseq data and performed computational biology. M.A. coordinated RNA sequencing strategy. Ca.J. provide guidelines for autophagy experiment design, LC3B IF protocol, and quantification of autophagic flux. S.R. provided HA15 compound (Cerezo et al. 2016). J.I and N.D provided PDX fragments and slices, PDX-derived cells and associated expertise. O.L. provided expertise on polysome-profiling in small tissue sample, translatome analysis and use of Anota2seq. S.S., S.P and Y.M. conceived the study. S.P., C.B. and Y.M. supervised the study and provided funding. S.S and Y.M. drafted the manuscript. R.N., Ch.J., S.P., C.B. edited the manuscript.

